# Social stress as a priming mechanism for Thermal Tolerance During a Poleward Range Shift

**DOI:** 10.1101/377283

**Authors:** Connor Wood, Robert N. L. Fitt, Lesley T. Lancaster

## Abstract

Cold tolerance plays a critical role in determining species’ geographical range limits. Previous studies have found that range shifts in response to climate warming are facilitated by cold acclimation capacities, due to increasingly colder and variable weather at high latitudes, and that cold tolerance can also be influenced by social factors. In this study we combined experiments and field studies to investigate the climatic and social factors affecting cold tolerances in range-shifting populations of the female-polymorphic damselfly *Ischnura elegans* in northeast Scotland. In the field, we observed both environmental (measured via habitat suitability) and social (sex ratio and density) effects on cold tolerance (CTmin). Androchrome females (male-like females) were less susceptible to beneficial social effects on cold tolerance than gynochromes (female-like females), and correspondingly, gynochrome frequency increased at colder, environmentally-limiting sites towards the range margin. Our manipulations of density in the laboratory further provide novel, experimental evidence that social interactions directly impact cold tolerance n this species. These results suggest that reciprocal effects of social environments on thermal acclimation may be an important but commonly overlooked aspect of allee effects which contribute to the formation of range margins. Moreover, our results point to a wider need to consider the role of population and social dynamics to shape both the thermal physiology of individuals and the thermal niches of species.

## Introduction

Understanding the mechanisms shaping thermal tolerance is critical for anticipating organismal responses to anthropogenic climate change. The 20th century has seen a rise in global average surface temperatures of about 0.6°C, with temperatures in the 21st century predicted to rise further, to somewhere between 0.6 – 4.0 °C (Field *et al*., 2014). Thermal tolerance plays an important role in determining a species’ fundamental niche, thus thermal tolerance may directly set a species’ geographic range limit (Brown *et al*., 1996; Wiens, 2011). As the climate warms and the geographical position of their thermal range limit changes with it, a species may expand its range to fill these new thermal boundaries. As a result, poleward shifts in species’ geographic ranges have already been documented in a wide range of species (Parmesan *et al*., 1999; Perry *et al*., 2005; Hickling *et al*., 2006; Chen *et al*., 2011; Mason *et al*., 2015).

Despite the fact that range shifts are facilitated by warmer climates, previous work suggests that species typically gain improved cold tolerance during shifts to higher latitudes and altitudes, putatively to facilitate survival in more variable, high-latitude climates (Lancaster *et al*., 2015; Lancaster, 2016). In fact, the capacity of a species to range shift in response to climate change may be limited by its ability to adapt or acclimate to novel cold-weather events beyond the historic range limit (Kellermann *et al*., 2009; Comte *et al*., 2014; Diamond, 2018). Ectotherms at an expanding northern range limit are therefore subject to intense selection pressure on their cold tolerances, and may commonly exhibit cold tolerances that are improved over cold tolerances observed in populations from the range core (Lancaster *et al*., 2015; Lancaster, 2016). Potentially by this process, terrestrial ectotherms commonly ‘overfill’ their range, and come to exist at latitudes beyond standard predictions of their cold tolerance as measured in the centre of the species’ range (Sunday *et al*., 2012). Understanding how range shifts both depend on and shape clinal variation in cold tolerance is therefore essential for understanding ongoing and future biodiversity shifts.

Cold tolerance clines can be created through natural selection or phenotypic plasticity, though typically by a combination of both (Schilthuizen & Kellermann, 2014). Numerous authors working on acclimation and adaptation have stressed the importance of considering phenotypic and developmental plasticity for range-shifting species (e.g. Nilsson-Ortman *et al*., 2012). In a model of adaptation, Lande (2009) suggested that extreme changes in environment can induce transient increases in genetic variance and phenotypic plasticity. This phase of increased adaptive potential, combined with the strong environmental effects on cold tolerance during poleward range-shifts, allows for rapid changes in phenotype and population structure. Under the Beneficial Acclimation Hypothesis (BAH) (Wilson & Franklin, 2002), and potentially due to genetic change, we should thus expect to find higher frequencies of cold tolerant phenotypes in marginal climates at the range edge of a range-expanding species.

On a physiological level, much of the adaptability in thermal tolerance can be attributed to the expression of heat shock proteins (HSPs; Dahlgaard *et al*., 1998; Zhao & Jones, 2012; Boykin *et al*., 2013; King & MacRae, 2015). Adaptive expression of HSPs is thought to part of a generalized stress response, allowing organisms to anticipate and respond to sudden extreme stressors (Nguyen *et al*., 2009; Benoit *et al*., 2010; Shu *et al*., 2011) as well as to adjust to the more regular stresses endured when living under non-optimal conditions (Sørensen *et al*., 2003). As a result of this, induction of HSPs by one stressor can have a ‘hardening’ effect which protects organisms from subsequent stresses.

There is growing awareness that various social stresses can also cause a generalized stress response that may carry over to affect individuals’ response to a range of other, non-social stressors (Pechenik, 2006; Sih, 2011). For example, Slos and Stoks (Slos & Stoks, 2008) were able to induce an increase in HSP70 in *Enallagma cyathigerum* larvae through non-lethal exposure to stickleback predators. In a similar study on *Coenagrion puella* larvae, Mikolajewski et al (2007) demonstrated that predator presence by fish, or cannibalistic pressure by conspecifics, reduced larval growth in both sexes. Moreover, in salmonid fish, dominance hierarchy formation induces HSP expression in both dominant and subordinate fish (Currie *et al*., 2010). There are very few studies, however, which have investigated directly whether intraspecific social stress can induce changes in thermal tolerance. In salmonid fish, social stress experienced by subordinate fish results in poorer cold tolerance, in comparison to dominant fish, despite their higher HSP expression (LeBlanc *et al*., 2011). In contrast, crowding stress in larval *Drosophila melanogaster* resulted in both increased HSP production and increased adult heat stress resistance (Sørensen & Loeschcke, 2001). These previous studies indicate that social stress can confer changes in thermal tolerance via a generalized stress response.

If social stress can induce either beneficial or detrimental effects on thermal tolerance, it could have major implications for population dynamics, particularly in range-expanding populations. Changes in social stress in range limit population may prime cold tolerance, facilitating colonization of cooler climates. We previously found that cold tolerance was positively associated with stressful social situations in range limit populations of the range-shifting damselfly *Ischnura elegans* (Lancaster *et al*., 2017a). In the current study, we expand on this finding in a new geographic region, and provide novel experimental evidence that social interactions shape beneficial thermal tolerances in a range-shifting species.

### Study system

The blue tailed damselfly *Ischnura elegans* (Vander Linden, 1820) is a small, coenagrionid damselfly that is ubiquitous throughout Europe and Central Asia. They are found in a wide variety of lowland water habitats, and have exhibited a recent range shifts into our study sites in the Scottish highlands (Cham *et al*., 2014), where the species has undertaken a range shift of 143 km northward during a recent, 60-year period; one of the largest recorded shifts in an Odonata species (Hickling *et al*., 2005). High-latitudes and elevations in Scotland are associated with highly variable climates and frequent, extreme-weather events in comparison to climates in central and southern Great Britain, and *I. elegans* has not yet been able to colonise high elevation sites within this region (Dijkstra & Lewington, 2006; Cham *et al*., 2014). Moreover, they remain at very low densities at higher latitudes (Hickling *et al*., 2005; Cham *et al*., 2014). Scottish populations of this species therefore present an ideal model system to investigate the drivers of novel cold tolerance phenotypes at the range margin and near the limits of their habitat suitability.

*Ischnura elegans* exhibits a genetically based, female-limited colour polymorphism: androchromes resemble males with blue-green thorax and blue abdominal segment 8, whereas infuscans (olive-green thorax, brown segment 8) and infuscans-obsoleta (brown thorax and segment 8) are visually distinct from males and are collectively referred to as gynochromes (Cordero *et al*., 1998). Site-specific colour morph frequencies have been shown to exhibit negative-frequency-dependent dynamics (Takahashi *et al*., 2014; Le Rouzic *et al*., 2015) because common morphs exhibit decreased fecundity due to mating harassment under their scramble competition mating system (Van Gossum *et al*., 2001; Gosden & Svensson, 2007; Sánchez-Guillén *et al*., 2017). Moreover, androchromes may receive additional protection from harassment due to their resemblance to males (intersexual mimicry (intersexual mimicry; (Gosden & Svensson, 2009). Male-male competition for food and mates can, in turn, be a powerful selective agent on male damselflies (Gosden & Svensson, 2008). Thus, social stress is a powerful evolutionary force which can cause rapid changes in both population size and allele frequencies in this species (Svensson *et al*., 2005; Gosden & Svensson, 2009).

We previously identified a latitudinal cline in cold tolerance in Swedish populations of *I. elegans* (Lancaster *et al*., 2015, 2016, 2017a; Dudaniec *et al*., 2018). Individuals from range edge populations had significantly faster chill coma recovery than core individuals. Chill coma recovery time was correlated with the minimum temperature experienced within the last 7 days, but only in range-limit populations (Lancaster *et al*., 2015). These results demonstrate adaptations of cold tolerance consistent with the BAH. Moreover, we also previously found that social feedback may play a critical role in the clinal variation of both cold tolerance and female colour morph frequencies for *I. elegans*. For androchromes, among-population variation in cold tolerance was best explained by recent cold weather events – a plastic response to thermal conditions consistent with the BAH. For gynochromes however, cold tolerance was positively correlated with gynochrome frequency in the population (Lancaster *et al*., 2017a). Moreover, the continent-wide latitudinal cline in morph frequencies for this species (increasing androchromes with latitude; Gosden *et al*., 2011) was found to be reversed at the range limit, where a high frequency of gynochromes was found at the most marginal or recently colonized sites (Lancaster *et al*., 2017a). This previous result suggests a beneficial feedback between gynochrome frequency, social stress, and cold tolerance that, under colder conditions, outweighed the typical frequency-dependent disadvantage of high gynochrome frequencies accruing from male-mating harassment. We previously suggested that the social stress involved with male-harassment may have had a cold tolerance priming effect in *I. elegans*, and as gynochromes lack the protection from male-mimicry and are thus particularly susceptible to higher harassment rates, they become subject to positive-frequency-dependent selection based on their more readily stress-induced phenotype at the colder range limit (Lancaster *et al*., 2017a).

In this study we investigated the alternative social and environmental mechanisms by which thermal tolerance is inducible in Scottish range shifting populations of *I. elegans*. Thermal tolerances were assessed at both larval and adult stages for wild-caught individuals from a geographically-dispersed range of study sites, each within 100km of the species’ elevational range margin, in order to examine the influence of environmental and social factors on cold tolerance across a range of social and environmental conditions. Further, social stress was generated via experimental social crowding of larvae in the laboratory, and thermal tolerances were then assessed to identify whether social crowding induced changes in cold tolerance in comparison to isolated individuals. We hypothesized that both colder climates and social crowding would increase the cold tolerances of damselflies during larval and adult stages. Based on our previous findings, we further expected that in females, gynochrome morphs would show a stronger relationship between social factors and cold tolerance than androchromes, and that this process may drive clinal variation in morph frequencies at the range limit.

## Materials and methods

### Social crowding study

A total of 161 individual *I. elegans* larvae were captured for the social crowding study, between the 5^th^ and 18^th^ of May 2017. Larvae were sampled from Midmar Stillwater Fishery (For site information, see Table 1, Figure 1). Larvae were collected from this single site in order to mitigate the confounding effects of climate differences among sites on larval physiology and thermal tolerances. Larvae were captured via dip netting and an initial identification was made using (Cham, 2012). *Ischnura elegans* larvae were transferred to plastic containers with air holes for transport back to the laboratory at the University of Aberdeen. Further identification and measurement were conducted using a Yenway SZN71 stereo microscope with a YenCam10 microscope camera (Ningbo Sunny Instruments Co. Ltd., Zhejiang, China), and images analysed with YenCam software (Version 3.6.9). Head width in mm (widest distance between the outer margins of the eyes) and number of caudal lamellae were recorded for each individual. All damselfly larvae have 3 caudal lamellae at birth, but some may be lost during development or capture. Caudal lamellae function as gills, therefore loss of lamellae may lower a damselfly larvae’s tolerance to heat and oxygen stress (Verberk & Calosi, 2012). Following the methodology of previous damselfly larvae research (e.g. Rolff, 1999), head widths were used as an indicator of larvae size, as head widths are well correlated with overall body size.

**Figure 1:**
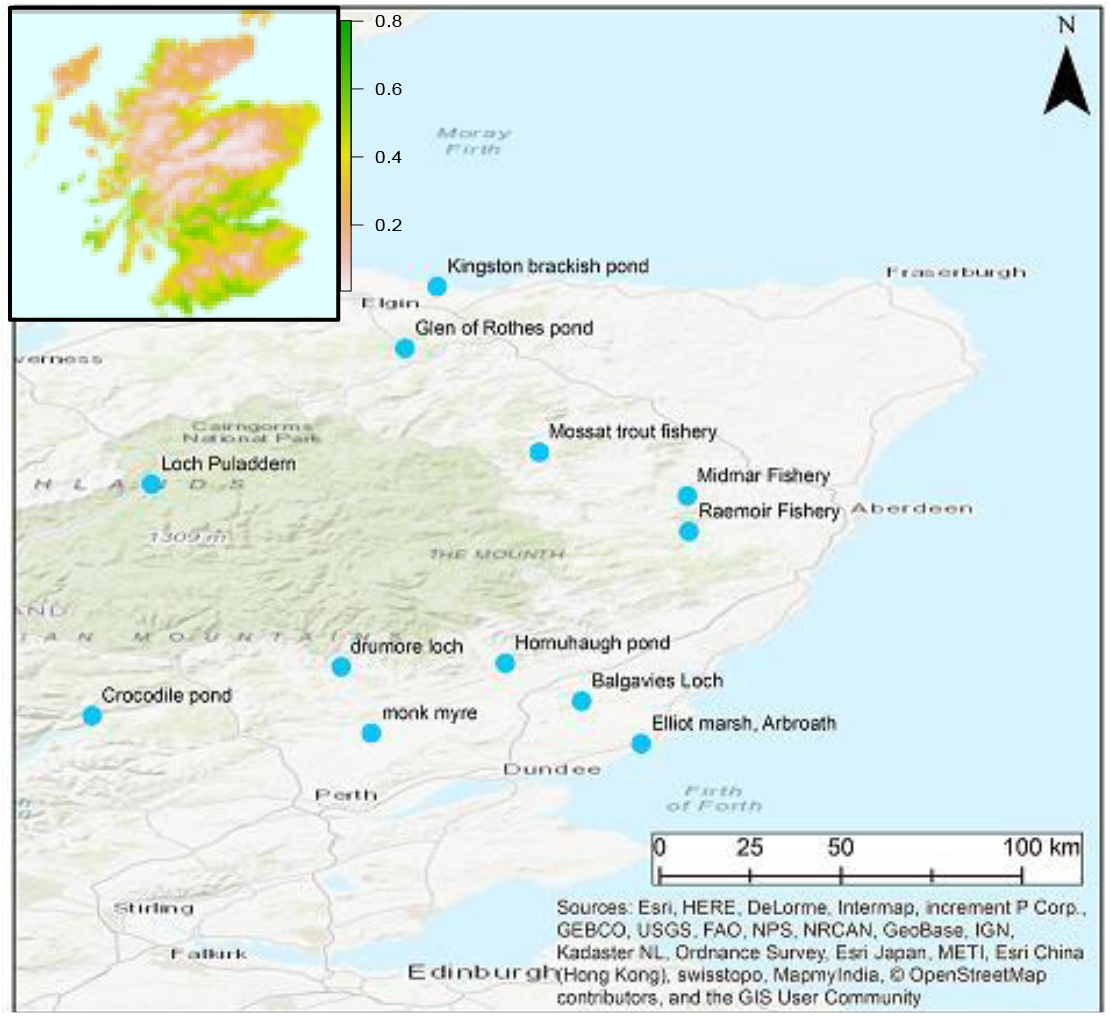
Locations of study sites near the elevational range limit for Ischnura elegans in Scotland. Inset shows habitat suitability for this species, with more suitable areas depicted in green (details of I. elegans habitat suitability estimation reported in (Fitt & Lancaster 2017). Map created using Arcmap (ESRI 2012; Redlands, CA, USA).

**Table 1:**
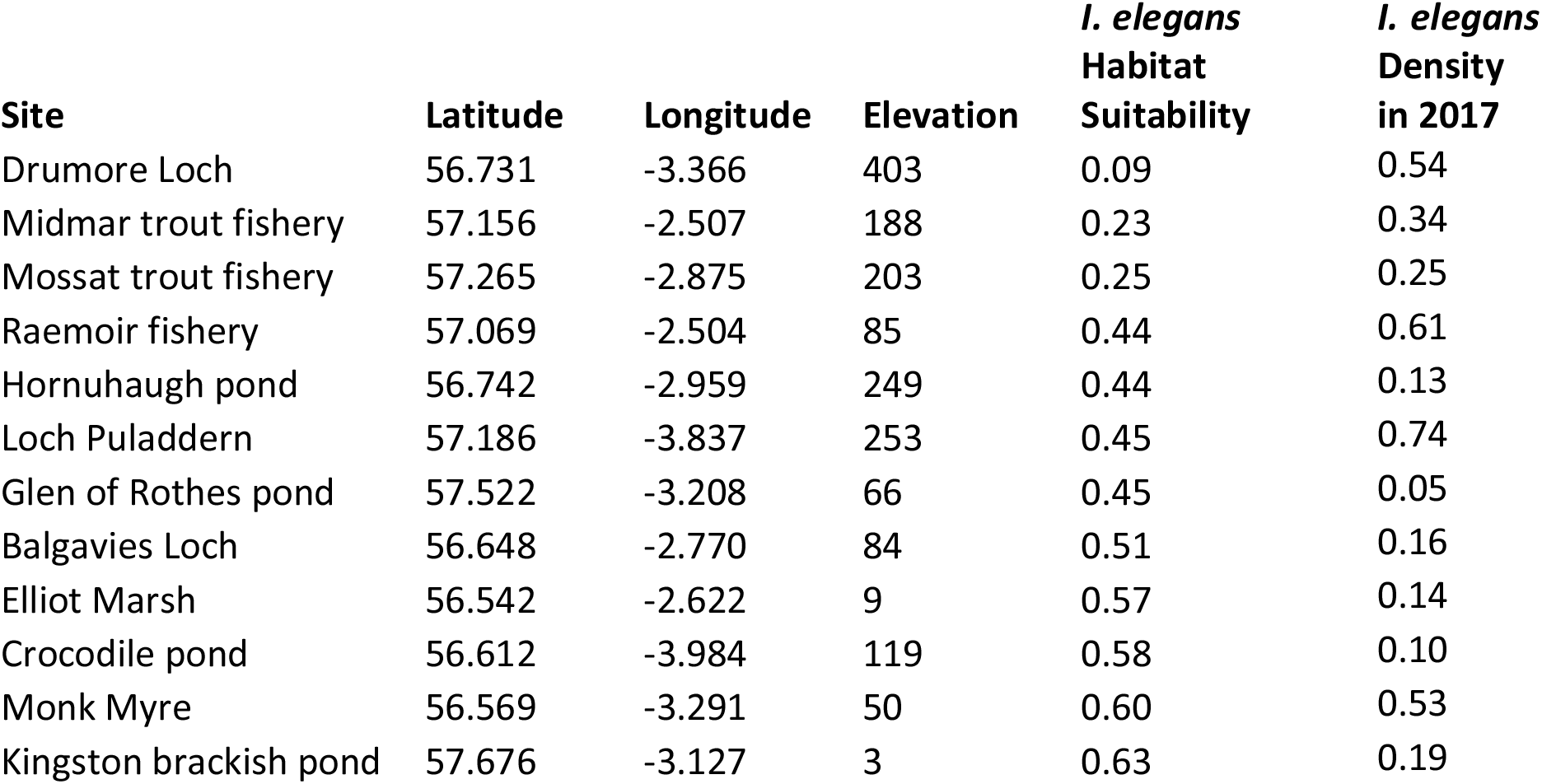
Location and Ischnura elegans habitat suitability values for our 12 study sites near the northern range limit in Scotland. Density estimates are individuals captured per minute.

| **Site** | **Latitude** | **Longitude** | **Elevation** | ***I. elegans***<br>**Habitat**<br>**Suitability** | ***I. elegans***<br>**Density**<br>**in 2017** |
| --- | --- | --- | --- | --- | --- |
| Drumore Loch | 56.731 | -3.366 | 403 | 0.09 | 0.54 |
| Midmar trout fishery | 57.156 | -2.507 | 188 | 0.23 | 0.34 |
| Mossat trout fishery | 57.265 | -2.875 | 203 | 0.25 | 0.25 |
| Raemoir fishery | 57.069 | -2.504 | 85 | 0.44 | 0.61 |
| Hornuhaugh pond | 56.742 | -2.959 | 249 | 0.44 | 0.13 |
| Loch Puladdern | 57.186 | -3.837 | 253 | 0.45 | 0.74 |
| Glen of Rothes pond | 57.522 | -3.208 | 66 | 0.45 | 0.05 |
| Balgavies Loch | 56.648 | -2.770 | 84 | 0.51 | 0.16 |
| Elliot Marsh | 56.542 | -2.622 | 9 | 0.57 | 0.14 |
| Crocodile pond | 56.612 | -3.984 | 119 | 0.58 | 0.10 |
| Monk Myre | 56.569 | -3.291 | 50 | 0.60 | 0.53 |
| Kingston brackish pond | 57.676 | -3.127 | 3 | 0.63 | 0.19 |

After collection and measurement, *I. elegans* larvae were randomly assigned to one of two social crowding treatments and were transferred to cylindrical enclosures (diameter 6.75 cm, height 3.5 cm) filled with 45-50 mL of 24+ hour aged tap water. The enclosures were sealed, although air holes in the lids permitted oxygen transfer. A fine mesh wall divided the enclosure into two compartments. Each compartment was roughly identical in size and contained a small section of plastic aquarium plant. Larvae in the ‘Isolated’ treatment were kept to a single compartment, while the other remained empty. In the Paired treatment, two larvae were transferred to the same enclosure, with one in each compartment. By separating individuals in the same waterbody with the mesh wall, the individual larvae were thus exposed to the visual- and chemical stimuli of their conspecifics in a contained environment whilst also being prevented from performing direct agonistic interactions such as cannibalism. Additionally, rearing the Isolated larvae in identical mesh enclosures ensured all larvae were exposed to the same physical environmental conditions during the study. Eighty-one larvae were randomly assigned to the Isolated treatment, and 80 to the Paired treatment.

Feeding occurred approximately every 24 hours, with larvae being fed 2-3 live aquatic invertebrates (e.g. brine shrimp *(Artemia)* bloodworms *(Glycera)*, and *Gammarus*). Water in each replicate was aerated during feeding, and approximately 20% water changes occurred every 5-7 days, or more frequently if required. Enclosures were maintained at 15 °C in a refrigerated incubator for the duration of the study, which approximates summer freshwater temperature in Scotland (Hrachowitz *et al*., 2010). The larvae were reared in these enclosures until the 19^th^ of June (32-45 days, depending upon capture date). At the end of the rearing treatment, 39 individuals from the Isolated treatment and 40 individuals from the Paired treatment survived for cold tolerance trials.

### Social and environmental determinants of cold tolerance in the wild

We sampled a total of 856 damselflies during June 2017, conducting repeat visits to 12 pond/loch sites throughout Northeast Scotland near the northern range limit for this species (Figure 1). The sites chosen were characterised as permanent water bodies with open, shallow bottoms and consistent varieties of emergent vegetation. They covered a large spatial extent and a range of climatic variation: Latitude ranges from 56.37° to 57.68° and longitude from −4.13° to −2.11°, with elevation ranges from 3 m to 442 m. The selected sites had previously been studied for their established *I. elegans* populations, with population density estimates available for the years 2014-2016 (data deposited in BioTIME: Dornelas *et al*., 2018). In total we sampled 551 *I. elegans* which we used to calculate sex ratio, population density, and female morph frequencies at each site. Across all sites, the 551 *I. elegans* comprised 247 males (44.82%), 196 females (35.57%) and 108 larvae (19.6%). Of these 551 individuals, we assessed cold tolerance for n = 375 *I. elegans*, including 160 adult females (n=82 androcromes, n= 36 gynochromes, n= 42 immature adult morphs), 107 adult males, and 108 larvae.

### Cold Tolerance Trials

Thermal tolerance for adult and larvae damselflies was measured as the critical thermal minimum (CTmin; i.e. the temperature at which individuals lost muscle control). CTmin was assessed using a thermal ramping protocol. Damselflies were placed individually in test tubes which were suspended in a Grant TX150 programmable circulating water bath with C2G cooling attachment (Grant Instruments, Shepreth, UK). Adults were placed in 50ml centrifuge tubes open to the air, and larvae in tubes filled with dechlorinated tap water at 15 °C. After a 10 minute equilibrium period at 15 °C, the water bath temperature was gradually decreased at a rate of 0.1° C min^−1^ until loss of movement and responsiveness was observed (individual no longer responded to prodding). Once CTmin had been recorded, damselflies were removed from the water bath, and individuals from both the wild and the rearing experiment were measured. Larval head widths were recorded as described above. Adult body length was assessed by scanning individuals on an Epson perfection photo scanner (Suwa, Nagano, Japan), and analyzing images with Fiji software (Schindelin *et al*., 2012).

### Data analysis

We analysed the relationships between CTmin and social/environmental factors for freshly-captured individuals across a range of sites, to test for the potential role of social interactions on thermal tolerance for *I. elegans* in the wild. Separate linear mixed models were run for adult and larval wild-caught damselflies, where each model included effects for individual traits, as well as social and environmental predictors. Social predictors included population density, sex ratio and colour morph frequencies. These were estimated for the *I. elegans* population at each site using the capture data in the current year (2017). Density estimates were calculated as the number of individuals captured per minute of catching effort (CPUE; Lancaster et al., 2015; Svensson et al., 2005). Sex ratio and morph frequency estimates were calculated under the assumption that catchability did not differ significantly between males, females and colour morphs (Stoks, 2001), thus CPUE was representative of the relative abundances of each sex/morph. Sex ratio was calculated as the proportion of males in the adult population per site. Colour-morph frequencies were estimated as the number of androchromes divided by the number of androchromes + gynochromes caught per site. Data on *I. elegans* site density estimates from 2016 were also included in models to test for effects of population densities during development on thermal tolerances (in the study region, *I. elegans* can take as long as 3 years to mature, with most likely maturing in 2 years; Fitt, Hand, and Lancaster, unpublished).

These social effects were contrasted with potential climatic effects on CTmin. To model climatic effects, we included site-specific values for habitat suitability, which was estimated using Maxent models for a previous study (Fitt & Lancaster, 2017), and including 9 non-correlated bioclimatic variables (Hijmans *et al*., 2005), of which bio1 (mean annual temperature) had the highest permutation importance (54%) followed by bio10 (mean temperature of the warmest quarter; 15.5%) and bio2 (mean diurnal temperature range; 11%). To identify recent acclimation effects on thermal tolerance, we also included an estimate of the minimum temperatures recorded in the region over the 6 days prior to capture (data from accuweather.com). In addition to social and environmental factors, we also included fixed effects for the individual traits body size (head width for larvae, body length for adults) and sex (adult analysis only).

We included all fixed terms and their interactions and performed AICc comparisons of different hypothesized combinations of social and environmental determinants of cold tolerance, to arrive at best-fit models. Analyses were performed in the lme4 and lmerTest packages for R v. 3.5.0 (R Core development Team, 2012; Bates *et al*., 2014; Kuznetsova *et al*., 2014). AICc was calculated using the AIccmodavg package for R (Mazerolle, 2015). Collinearity of fixed effects was assessed using the vif.mer() function for R (Frank, 2014). Site ID and capture date were included as random factors in all models. CTmin was log-transformed (log[CTmin+2]) to improve normality and homogeneity of the residuals for the models. For adult females, we also conducted a targeted hypothesis test to identify if any of our social or environmental predictors affected cold tolerance differentially among female morphs. For this, mixed models were run as described above, but the analysis was limited to adult females only. We examined each social and environmental factor for significant interactions with female morphotypes. Non-significant interactions (and their associated main effects) were sequentially removed from the analysis.

Using data on morph frequencies, population densities and sex ratios gathered in 2014 from n=93 sites across the study region (Fitt & Lancaster, 2017; Dornelas *et al*., 2018), we also examined relationships between social and environmental predictor variables using linear models in R, and including latitude and longitude to investigate spatial trends.

For the social crowding study, we analysed the drivers of CTmin using linear mixed models as described above, in which treatment (Paired vs. Individual), head width, numbers of caudal lamellae (1-3) and all interactions were included as fixed effects. A random effect for container was also included in the models. To assess the potential effects of hypoxia on thermal tolerance, we analysed existing data from n=20 damselflies from a previous study which used the same rearing protocol described here, but which explicitly manipulated oxygenation levels (Morrison & Lancaster, unpublished).

## Results

Across sites, CTmin of adult male, adult female, and larval *I. elegans* captured from the field were: 3.92 ± 0.32 (SE) °C, 3.15 ± 0.26 °C, and 2.13 ± 0.22 °C respectively. The best model explaining cold tolerance of wild-caught, adult damselflies from our 12 sites (Table 2A) included a significant 3-way interaction among sex, habitat suitability, and sex ratio and a 2-way interaction between sex and site density in the current year. Male cold tolerance was best explained by the interaction between habitat suitability and population sex ratio, such that males from poor-quality sites had lowest (best) cold tolerance overall, but cold tolerance was also lower (better) at high quality sites when the proportion of males in the population was high (Figure 2A; effect of habitat suitability * sex ratio in males only = −18.82±4.72, t=−4.21, P=0.006, this effect is n.s. in females). In contrast, female cold tolerance was best explained by population density, such that female cold tolerance improved (was lower) with higher population density (Figure 2D; effect of density in females = −0.69±0.22, t=−3.19, P=0.002, this effect is n.s. in males) and with lower habitat suitability (Figure 2B; effect of habitat suitability in females = 1.88±0.41, t=4.64, P=0.0005). No other trait or social or environmental effect significantly affected cold tolerance.

**Figure 2:**
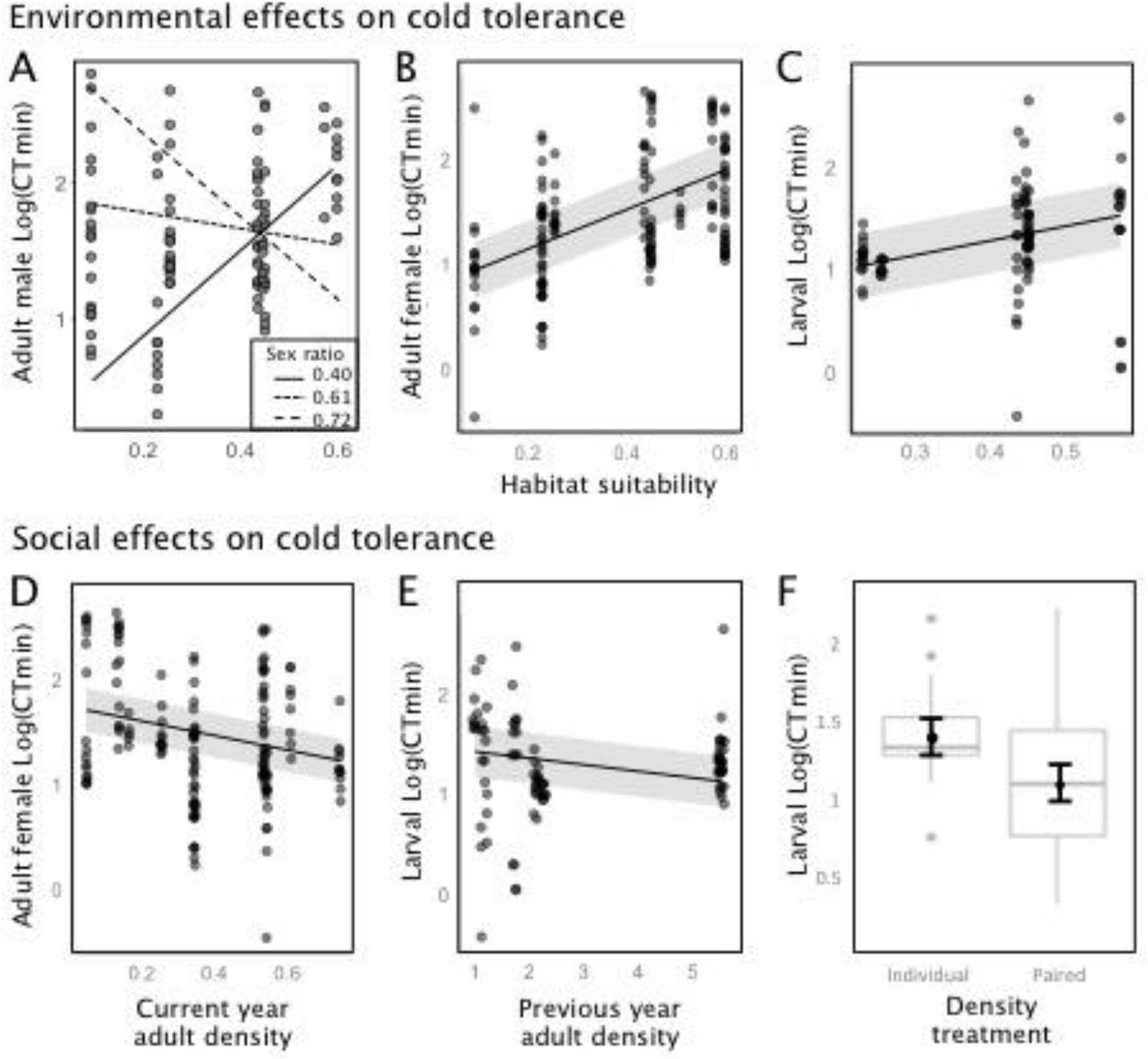
Environmental (A-C) and social (D-F) determinants of cold tolerance in adult and larval Ischnura elegans captured near their elevational range limit is Scotland.

**Table 2:**
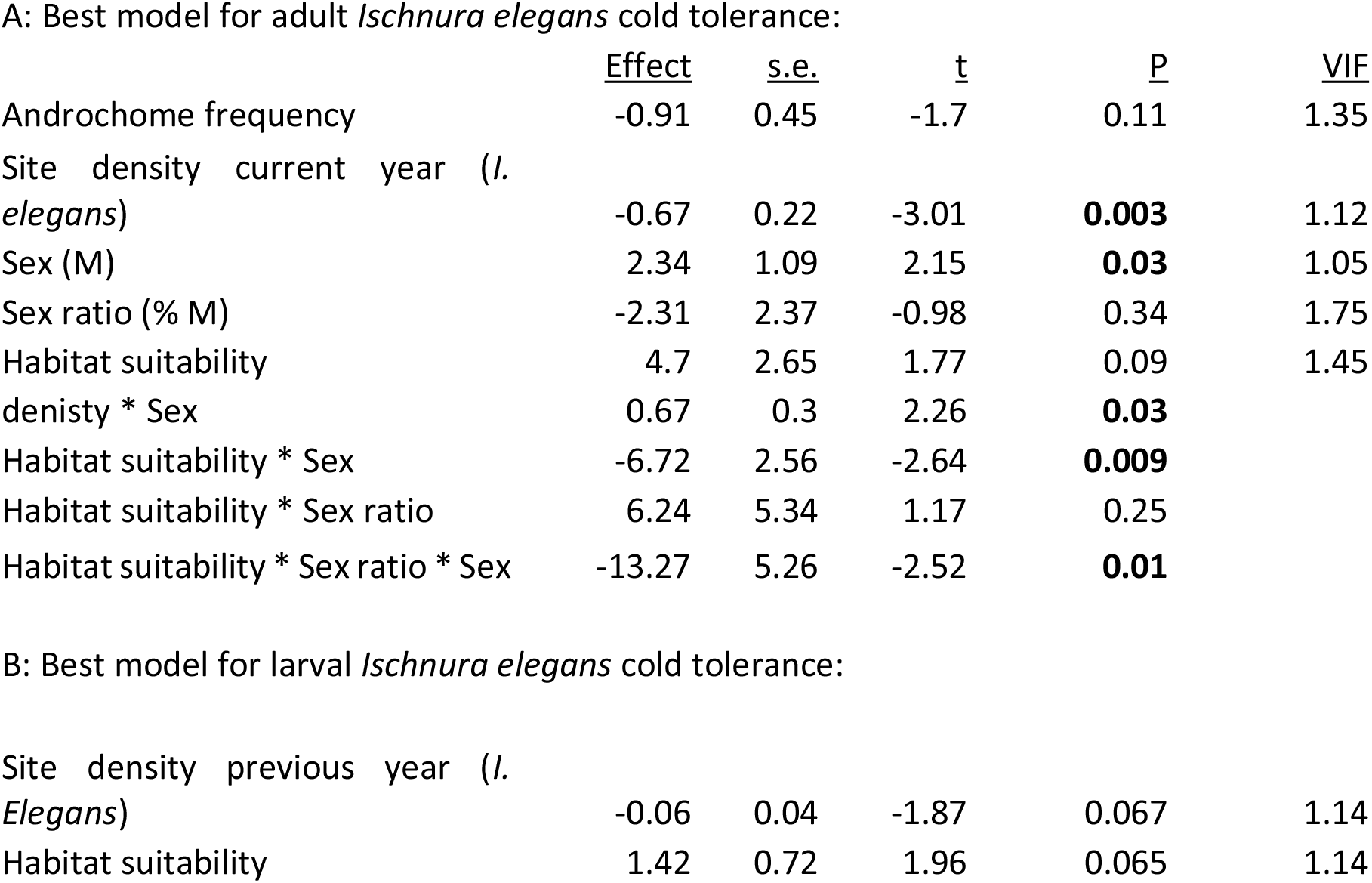
Fixed-effects in best-fit models for adult (A) and larval (B) Ischnura elegans cold tolerances, in wild-caught individuals from our 12 range limit populations.

A: Best model for adult *Ischnura elegans* cold tolerance:
|  | <u>Effect</u> | <u>s.e.</u> | <u>t</u> | <u>P</u> | <u>VIF</u> |
| --- | --- | --- | --- | --- | --- |
| Androchome frequency | -0.91 | 0.45 | -1.7 | 0.11 | 1.35 |
| Site density current year (*I. elegans*) | -0.67 | 0.22 | -3.01 | **0.003** | 1.12 |
| Sex (M) | 2.34 | 1.09 | 2.15 | **0.03** | 1.05 |
| Sex ratio (% M) | -2.31 | 2.37 | -0.98 | 0.34 | 1.75 |
| Habitat suitability | 4.7 | 2.65 | 1.77 | 0.09 | 1.45 |
| denisty * Sex | 0.67 | 0.3 | 2.26 | **0.03** |  |
| Habitat suitability * Sex | -6.72 | 2.56 | -2.64 | **0.009** |  |
| Habitat suitability * Sex ratio | 6.24 | 5.34 | 1.17 | 0.25 |  |
| Habitat suitability * Sex ratio * Sex | -13.27 | 5.26 | -2.52 | **0.01** |  |
| B: Best model for larval *Ischnura elegans* cold tolerance: |  |  |  |  |  |
| Site density previous year (*I. Elegans*) | -0.06 | 0.04 | -1.87 | 0.067 | 1.14 |
| Habitat suitability | 1.42 | 0.72 | 1.96 | 0.065 | 1.14 |

Corroborating previous findings in Swedish populations of *I. elegans*, we found that gynochrome females exhibited stronger social controls on cold tolerance than androchromes, such that gynochrome females exhibited improved cold tolerance after having developed in high density sites, whereas androchromes were unaffected by developmental densities (effect of previous year population density * female morph = −0.11±0.04, t=−2.99, P=0.003; Figure 3A). No other social or environmental factor interacted with female morphotype to affect cold tolerance, and, as in Swedish populations, there were no clear, absolute differences in the cold tolerances of these alternative morphotypes, suggesting that the alternative morphotypes differ in drivers of acclimation processes but not in their absolute capacities to express cold tolerance.

**Figure 3:**
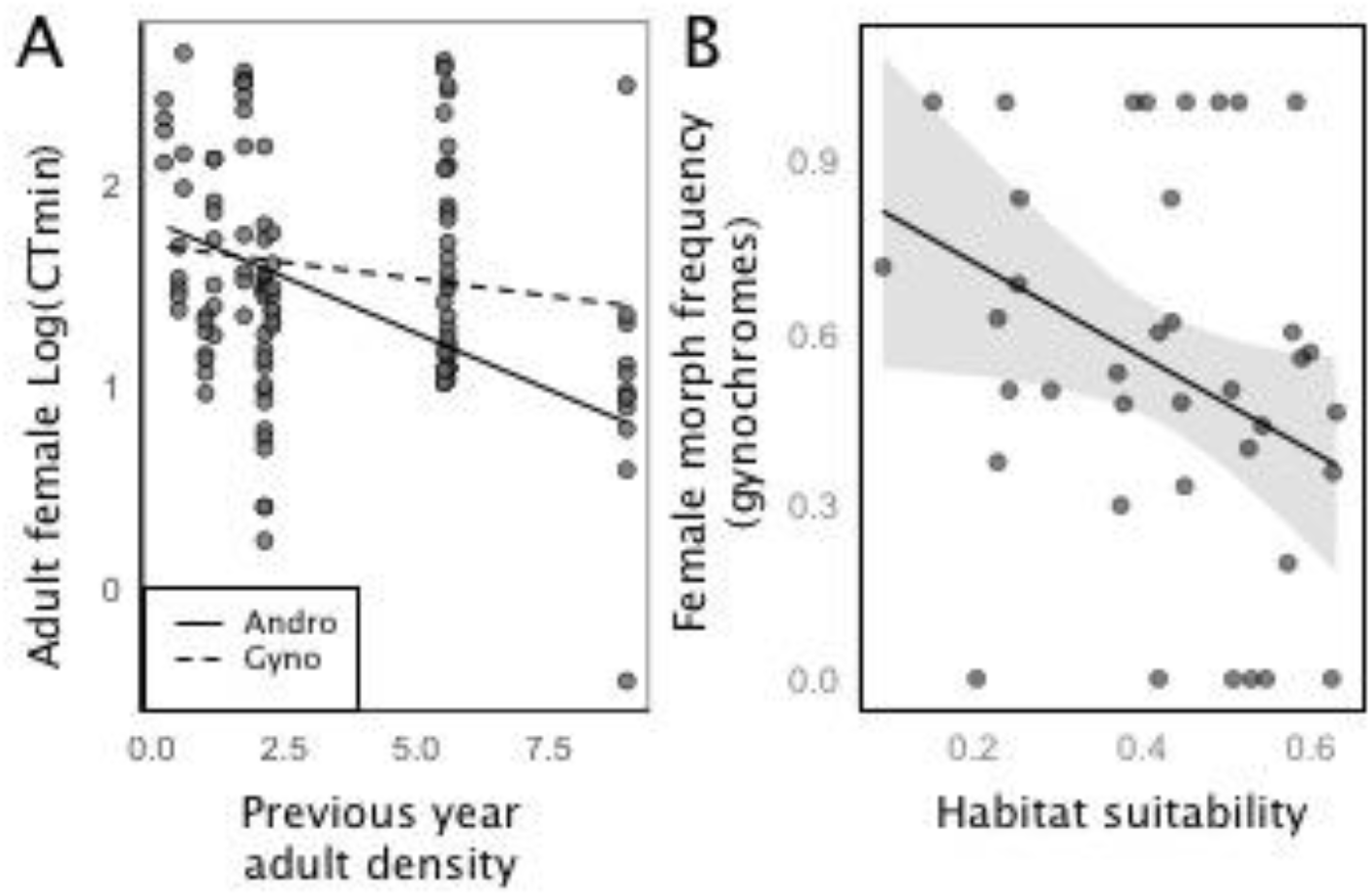
A) Differential effects of developmental densities on adult thermal tolerances by morphotype; gynochromes are more likely to experience beneficial social priming. B) Gynochromes are more frequent in lower-suitability sites near the elevational range margin, suggesting that social priming effects on gynochrome cold tolerance are adaptive there.

Cold tolerances of wild-caught larvae were best explained by a model which included habitat suitability and population density in the previous year (Table 2B). As in adults, larval cold tolerance was lower (better) in sites characterized by lower habitat suitability (Figure 2C). Wild-caught larval thermal tolerance was also improved (lower) in sites characterized by high developmental densities (previous year adult densities; Figure 2E), corroborating our experimental findings (see below) that larval competition has a beneficial effect on cold tolerance. No other social or environmental factor influenced larval cold tolerances. Body size did not significantly affect cold tolerance in either adults or larvae.

As in (Lancaster et al. 2017), we found that gynochrome frequencies were higher in cooler, less suitable habitats among range limit sites (effect of habitat suitability on gynochrome frequency = −0.81±0.39, t=−2.10, P=0.04; Figure 3B). Also consistent with (Lancaster *et al*., 2015), we find that Scottish populations of *I. elegans* tend to achieve higher population densities at more suitable sites (effect of habitat suitability on log(density) = 0.27±0.12, t=2.33, P=0.02).

The best model for cold tolerance of larvae from our rearing experiment included only a single fixed effect for treatment (effect of pairing vs. isolation = −0.27±0.08, t=−3.40, P=0.001). Individuals reared in the paired treatment experienced significantly improved (lower) cold tolerance in comparison to individually-housed larvae (Figure 2F; individual larvae expressed an average CTmin of 2.23 ± 0.19(SE) °C, while paired larvae expressed a CTmin of 1.40 ± 0.24 °C). Effects of body size and number of tail segments were not significant and worsened model fit. There was no effect of dissolved oxygen levels within containers on the thermal tolerance of larvae (effect of dissolved O2 concentration on cold tolerance = −0.92±1.43 (s.e.), t=−0.64, P=0.53). Additionally, for individuals in the Paired treatment, relative- and absolute difference in head width between paired conspecifics were found to have no significant effect on CTmin. There was also no significant difference between treatments in head width, number of lamellae, growth rate or treatment duration (p = 0.489, 0.78, 0.328 and 0.07 for head width, lamellae, growth rate and treatment duration, respectively).

## Discussion

We investigated social and environmental drivers of cold tolerance in marginal populations of a range-shifting damselfly, addressing the hypothesis that social factors can influence colonization of stressful environments. Based on our previous findings, we posited that social interactions can initiate a generalised stress response that offers protection against novel climatic variation. As expected, and in accordance with the Beneficial Acclimation Hypothesis, all individuals exhibited better tolerance to cold when they were captured from sites characterized by lower climatic habitat suitability (i.e., generally cooler and more climatically variable sites, see methods and Fitt & Lancaster, 2017; Figure 2A-C). This effect is likely to reflect the species’ capacity for phenotypic plasticity (Lancaster *et al*., 2017b), but may also contain some element of local adaptation in cold tolerances. A recent population genomic study of these populations revealed generally weak genetic structure and low genetic divergence across the region, and also suggested high levels of gene flow among most sites (Fitt and Lancaster, unpublished data). Specifically, significant genetic differentiation was observed at a few isolated, high elevation sites (notably Loch Puladdern and Honuhaugh Pond; Figure 1), sites which were also associated with atypical values for climatic variability. Moreover, in a separate region, we found significant genetic differentiation in genetic variants mapping to HSP70 across the climatic gradient over which *I. elegans* has recently expanded (Dudaniec et al., 2018), as well as latitudinal variation in cold tolerance plasticity (Lancaster *et al*., 2015). More work is needed to disentangle the genetic and plastic components of the effect of habitat suitability on cold tolerance of these range shifting damselflies.

Interestingly, we also found evidence for significant social influences of the expression of cold tolerance in larvae and adults of both sexes within range-limit populations of this species (Figure 2D-F). For adult male *I. elegans*, cold tolerance was best explained by an interaction between sex ratio and habitat suitability, such that males typically exhibited better tolerance to cold either following direct acclimation to cooler climates (see above). However, in the absence of direct acclimation to cold weather, they could also achieve improved cold tolerance when at high frequency in the population. Males are likely to experience social stress at high frequency due to increasingly intense competition for mates (Cordero *et al*., 1997), or due to other forms of male-male interference for feeding or basking sites (Baird & May, 2003; Fitt & Lancaster, 2017), and this source of social stress evidently is associated also with increased resistance to cold. In contrast to males, females and larvae both experienced increased cold tolerance in response to high population density. Social stress associated with high densities likely reflects both increased predation pressure and competition for food, basking, and oviposition sites (Pechenik, 2006). For adult females, high densities also reflect more intense mating harassment overall. Van Gossum et al. (2001) previously found that mating harassment rates, and particularly the tendency of males to target common female morphotypes, increased at high population densities, suggesting that common morphs will be most prone to experiencing the cold tolerance benefit of density-induced social stress.

Moreover, the current study demonstrates experimentally that crowding has a direct, beneficial effect on cold tolerance expression in larval damselflies. As our study design prohibited the role of agonistic interactions by separating the individuals into compartments, we can infer that the effect on cold tolerance was caused by indirect stress due to visual or chemical cues. Previous studies have demonstrated that damselfly physiological stress levels and HSP production are mediated by visual cues of conspecifics and predators (McPeek *et al*., 2001; Slos & Stoks, 2008). Analysis of the oxygenation dataset confirmed that hypoxia has no significant effect on cold tolerance for *I. elegans* larvae. These findings are consistent with previous research (Boardman *et al*., 2016). While some studies have demonstrated that anoxia can induce cold-hardening (in flies (in flies; (Coulson & Bale, 1991; Nilson *et al*., 2006), even if this were the case for *I. elegans*, the water in each enclosure was aerated frequently enough to discount anoxia as an influential factor. The increased cold tolerance for the paired larvae can therefore be best explained as a physiological response to chronic social stress.

As in Sweden (Lancaster *et al*., 2017a), we found that the continental-wide cline of increasing androchrome frequency with latitude was reversed at the northern range limit in Scotland (Figure 3B). Scottish range limit sites characterized by the lowest habitat suitabilities were dominated by the gynochrome morph. Moreover, as in Swedish range limit females, we find that Scottish range limit gynochromes are more susceptible than androchromes to beneficial effects of social stress on the expression of their cold tolerances. In the current study we find that gynochrome females who experienced high density during development were more likely to exhibit improved cold tolerance as adults, whereas androchromes were less likely to receive this social benefit (Figure 3A). If social stress is an important source of phenotypic variation in cold toelrance, the differences in social sensitivity among these morphs may explain why gynochomes reach high frequency at the range margin.

Although populations were likely to reach higher density at greater levels of habitat suitability (i.e., further from the range margin), all of our sites are within 100km of the range margin in Scotland, and high gene flow across the region suggests that this species is not dispersal limited across all of our study sites (Fitt and Lancaster, unpublished data). Therefore, populations persisting at the lowest levels of habitat suitability are likely to represent sink habitats which are consistently replenished by individuals from more suitable sites. The results presented here suggest that the survival of immigrants is likely to be positively-density-dependent, where relatively larger colonising population sizes confer a survival advantage to cold weather events; gynochromes are more likely to reap this social benefit and thus achieve higher frequencies in these, otherwise sink, habitats.

The observed requirement for social dynamics to augment cold tolerance at the range margin supports the idea that species’ thermal acclimation potential is not always sufficient alone to cope with novel climatic variation (Gunderson & Stillman, 2015; Lancaster *et al*., 2017a). Moreover, thermal acclimation benefits of social interactions reported here may be an important but commonly overlooked aspect of allee effects which contribute to the formation of range margins (Sexton *et al*., 2009). These results also highlight the need for further research into the extent to which social or sexual systems and group dynamics contribute to the thermal physiology of individuals, and the thermal niches of species.

## Author contributions

CW conceived the idea for the study in discussion with LL. CW and LL developed the experimental design and conducted the fieldwork. CW conducted the experiments. CW and LL analysed the data. CW drafted the ms. RF contributed to previous years’ data collection and generating the habitat suitability layer. All three authors contributed critically to the drafts and gave final approval for publication.

## Acknowledgements

We thank Mike Hinchliffe, Thomas Price, and Katie Grimmond for their assistance with fieldwork. We further thank all landowners for permission to work on their land, and particularly wish to acknowledge the proprietors of Midmar Trout Fishery for their hospitality. Erik I. Svensson provided thoughtful comments on a previous draft of this ms. This project was funded by the University of Aberdeen School of Biological Sciences MSc Ecology and Conservation programme.

## Data accessibility

All experimental and field data associated with this study will be deposited with datadryad.org and in BioTIME upon publication, and is available upon request from the authors.

## References

Baird, J.M. & May, M.L. 2003. Fights at the Dinner Table: Agonistic Behavior in Pachydiplax longipennis (Odonata: Libellulidae) at Feeding Sites. J. Insect Behav. 16: 189–216. Kluwer Academic Publishers-Plenum Publishers.

Bates, D., Mächler, M., Bolker, B. & Walker, S. 2014. Fitting Linear Mixed-Effects Models using lme4. J. Stat. Softw. 67: 51.

Benoit, J.B., Lopez-Martinez, G., Phillips, Z.P., Patrick, K.R. & Denlinger, D.L. 2010. Heat shock proteins contribute to mosquito dehydration tolerance. J. Insect Physiol. 56: 151–156.

Boardman, L., Sørensen, J.G., Koštál, V., Šimek, P. & Terblanche, J.S. 2016. Cold tolerance is unaffected by oxygen availability despite changes in anaerobic metabolism. Sci. Rep. 6: 32856. Nature Publishing Group.

Boykin, L.M., Bell, C.D., Evans, G., Small, I. & De Barro, P.J. 2013. Is agriculture driving the diversification of the Bemisia tabaci species complex (Hemiptera: Sternorrhyncha: Aleyrodidae)?: Dating, diversification and biogeographic evidence revealed. BMC Evol. Biol. 13: 228. BioMed Central.

Brown, J.H., Stevens, G.C. & Kaufman, D.M. 1996. The geographic range: size, shape, and internal structure. Annu. Rev. Ecol. Syst. 27: 597–623.

Cham, S.A. 2012. Field guide to the larvae and exuviae of British dragonflies: damselflies (Zygoptera) and dragonflies (Anisoptera). British Dragonfly Society.

Cham, S., Nelson, B., Parr, A., Prentice, S., Smallshire, D. & Taylor, P. 2014. Atlas of Dragonflies in Britain and Ireland. the Field Studies Council for the Biological Records Centre, Centre for Ecology & Hydrology, with the British Dragonfly Society.

Chen, I.-C., Hill, J.K., Ohlemüller, R., Roy, D.B. & Thomas, C.D. 2011. Rapid range shifts of species associated with high levels of climate warming. Science 333: 1024–1026.

Comte, L., Murienne, J. & Grenouillet, G. 2014. Species traits and phylogenetic conservatism of climate-induced range shifts in stream fishes. Nat. Commun. 5: 5023. Nature Publishing Group.

Cordero, A., Carbone, S.S. & Utzeri, C. 1997. Male mating success in a natural population of Ischnura elegans (Vander Linden) (Odonata: Coenagrionidae). Odonatologica 26: 459–465. S.I.O.

Cordero, A., Carbone, S.S. & Utzeri, C. 1998. Mating opportunities and mating costs are reduced in androchrome female damselflies,Ischnura elegans(Odonata). Anim. Behav. 55: 185–197. Academic Press.

Coulson, S.J. & Bale, J.S. 1991. Anoxia induces rapid cold hardening in the housefly Musca domestica (Diptera: Muscidae). J. Insect Physiol. 37: 497–501. Pergamon.

Currie, S., LeBlanc, S., Watters, M.A. & Gilmour, K.M. 2010. Agonistic encounters and cellular angst: social interactions induce heat shock proteins in juvenile salmonid fish. Proc. Biol. Sci. 277: 905–13. The Royal Society.

Dahlgaard, J., Loeschcke, V., Michalak, P. & Justesen, J. 1998. Induced thermotolerance and associated expression of the heat-shock protein Hsp70 in adult Drosophila melanogaster. Funct. Ecol. 12: 786–793. Wiley/Blackwell (10.1111).

Diamond, S.E. 2018. Contemporary climate-driven range shifts: Putting evolution back on the table. Funct. Ecol., doi: 10.1111/1365-2435.13095. Wiley/Blackwell (10.1111).

Dijkstra, K.-D.B. & Lewington, R. 2006. Field Guide to the Dragonflies of Britain and Europe. British Wildlife Publishing, Gillingham, Dorset, UK.

Dornelas, M., Antão, L.H., Moyes, F. & Al., E. 2018. BioTIME: A database of biodiversity time series for the Anthropocene. Glob. Ecol. Biogeogr. 0: 1–26; https://doi.org/10.1111/geb.12729.

Dudaniec, R.Y., Yong, C.J., Lancaster, L.T., Svensson, E.I. & Hansson, B. 2018. Signatures of local adaptation along environmental gradients in a range-expanding damselfly (Ischnura elegans). Mol. Ecol. 27: 2576–2593.

Field, C.B., Barros, V.R. & Intergovernmental Panel on Climate Change. Working Group II. 2014. Climate change 2014: impacts, adaptation, and vulnerability: Working Group II contribution to the fifth assessment report of the Intergovernmental Panel on Climate Change. Cambridge University Press.

Fitt, R.N.L. & Lancaster, L.T. 2017. Range shifting species reduce phylogenetic diversity in high latitude communities via competition. J. Anim. Ecol. 86.

Frank, A.F. 2014. mer-utils.r. Github repository.

Gosden, T.P., Stoks, R. & Svensson, E.I. 2011. Range limits, large-scale biogeographic variation, and localized evolutionary dynamics in a polymorphic damselfly. Biol. J. Linn. Soc. 102: 775–785. Blackwell Publishing Ltd.

Gosden, T.P. & Svensson, E.I. 2009. Density-Dependent Male Mating Harassment, Female Resistance, and Male Mimicry. Am. Nat. 173: 709–721.

Gosden, T.P. & Svensson, E.I. 2007. Female Sexual Polymorphism and Fecundity Consequences of Male Mating Harassment in the Wild. PLoS One 2: e580.

Gosden, T.P. & Svensson, E.I. 2008. Spatial and temporal dynamics in a sexual selection mosaic. Evolution (N. Y). 62: 845–856. Wiley/Blackwell (10.1111).

Gunderson, A.R. & Stillman, J.H. 2015. Plasticity in thermal tolerance has limited potential to buffer ectotherms from global warming. Proc. Biol. Sci. 282: 20150401.

Hickling, R., Roy, D.B., Hill, J.K., Fox, R. & Thomas, C.D. 2006. The distributions of a wide range of taxonomic groups are expanding polewards. Glob. Chang. Biol. 12: 450–455.

Hickling, R., Roy, D.B., Hill, J.K. & Thomas, C.D. 2005. A northward shift of range margins in British Odonata. Glob. Chang. Biol. 11: 502–506.

Hijmans, R.J., Cameron, S.E., Parra, J.L., Jones, P.G. & Jarvis, A. 2005. Very high resolution interpolated climate surfaces for global land areas. Int. J. Climatol. 25: 1965–1978.

Hrachowitz, M., Soulsby, C., Imholt, C., Malcolm, I.A. & Tetzlaff, D. 2010. Thermal regimes in a large upland salmon river: a simple model to identify the influence of landscape controls and climate change on maximum temperatures. Hydrol. Process. 24: 3374–3391.

Kellermann, V., van Heerwaarden, B., Sgrò, C.M. & Hoffmann, A.A. 2009. Fundamental evolutionary limits in ecological traits drive Drosophila species distributions. Science 325: 1244–6. American Association for the Advancement of Science.

King, A.M. & MacRae, T.H. 2015. Insect heat shock proteins during stress and diapause. Annu. Rev. Entomol. 60: 59–75.

Kuznetsova, A., Brockhoff, P.B. & Christensen, R.H.B. 2014. lmerTest: Tests for random and fixed effects for linear mixed effects models (lmer objects of lme4package). R package version 2.0-1.1.

Lancaster, L.T. 2016. Widespread range expansions shape latitudinal variation in insect thermal limits. Nat. Clim. Chang. 6: 618–621.

Lancaster, L.T., Dudaniec, R., Chauhan, P., Wellenreuther, M., Svensson, E.I. & Hansson, B. 2016. Gene expression under thermal stress varies across a geographic range expansion front. Mol. Ecol. 25: 1141–56.

Lancaster, L.T., Dudaniec, R.Y., Hansson, B. & Svensson, E.I. 2017a. Do group dynamics affect colour morph clines during a range shift? J. Evol. Biol. 30.

Lancaster, L.T., Dudaniec, R.Y., Hansson, B. & Svensson, E.I. 2015. Latitudinal shift in thermal niche breadth results from thermal release during a climate-mediated range expansion. J. Biogeogr. 42.

Lancaster, L.T., Morrison, G. & Fitt, R.N. 2017b. Life history trade-offs, the intensity of competition, and coexistence in novel and evolving communities under climate change. Philos. Trans. R. Soc. B Biol. Sci. 372.

Lande, R. 2009. Adaptation to an extraordinary environment by evolution of phenotypic plasticity and genetic assimilation. J. Evol. Biol. 22: 1435–46.

Le Rouzic, A., Hansen, T.F., Gosden, T.P. & Svensson, E.I. 2015. Evolutionary time-series analysis reveals the signature of frequency-dependent selection on a female mating polymorphism. Am. Nat. 185: E182–96. University of Chicago PressChicago, IL.

LeBlanc, S., Middleton, S., Gilmour, K.M. & Currie, S. 2011. Chronic social stress impairs thermal tolerance in the rainbow trout (Oncorhynchus mykiss). J. Exp. Biol. 214: 1721–31.

Mason, S.C., Palmer, G., Fox, R., Gillings, S., Hill, J.K., Thomas, C.D., et al. 2015. Geographical range margins of many taxonomic groups continue to shift polewards. Biol. J. Linn. Soc. 115:586–597.

Mazerolle, M.J. 2015. AICcmodavg: Model selection and multimodel inference based on (Q)AIC(c). R package version 2.0-3.

McPeek, M.A., Grace, M. & Richardson, J.M.L. 2001. Physiological and behavioural responses to predators shape the growth/predation risk trade-off in damselflies. Ecology 82: 1535–1545. Wiley-Blackwell.

Mikolajewski, D.J., Stoks, R., Rolff, J. & Joop, G. 2007. Predators and cannibals modulate sex-specific plasticity in life-history and immune traits. Funct. Ecol. 0: 071107024542001–??? Wiley/Blackwell (10.1111).

Nguyen, T.T.A., Michaud, D. & Cloutier, C. 2009. A proteomic analysis of the aphid Macrosiphum euphorbiae under heat and radiation stress. Insect Biochem. Mol. Biol. 39: 20–30.

Nilson, T.L., Sinclair, B.J. & Roberts, S.P. 2006. The effects of carbon dioxide anesthesia and anoxia on rapid cold-hardening and chill coma recovery in Drosophila melanogaster. J. Insect Physiol. 52: 1027–33. NIH Public Access.

Nilsson-Ortman, V., Stoks, R., De Block, M. & Johansson, F. 2012. Generalists and specialists along a latitudinal transect: patterns of thermal adaptation in six species of damselflies. Ecology 93: 1340–52.

Parmesan, C., Ryrholm, N., Stefanescu, C., Hill, J., Thomas, C., Descimon, H., et al. 1999. Poleward shifts in geographical ranges of butterfly species associated with regional warming. Nature 399: 579–583. MACMILLAN MAGAZINES LTD, PORTERS SOUTH, 4 CRINAN ST, LONDON N1 9XW, ENGLAND.

Pechenik, J.A. 2006. Larval experience and latent effects--metamorphosis is not a new beginning. Integr. Comp. Biol. 46: 323–333. Oxford University Press.

Perry, A.L., Low, P.J., Ellis, J.R. & Reynolds, J.D. 2005. Climate Change and Distribution Shifts in Marine Fishes. Science (80-.). 308: 1912–1915.

R Core development Team. 2012. R: A language and environment for statistical computing. R foundation for Statistical Computing, Vienna, Austria.

Rolff, J. 1999. Parasitism increases offspring size in a damselfly: experimental evidence for parasite-mediated maternal effects. Anim. Behav. 58: 1105–1108. Academic Press.

Sánchez-Guillén, R.A., Wellenreuther, M., Chávez-Ríos, J.R., Beatty, C.D., Rivas-Torres, A., Velasquez-Velez, M., et al. 2017. Alternative reproductive strategies and the maintenance of female color polymorphism in damselflies. Ecol. Evol. 7: 5592–5602. Wiley-Blackwell.

Schilthuizen, M. & Kellermann, V. 2014. Contemporary climate change and terrestrial invertebrates: evolutionary versus plastic changes. Evol. Appl. 7: 56–67. Wiley/Blackwell (10.1111).

Schindelin, J., Arganda-Carreras, I., Frise, E., Kaynig, V., Longair, M., Pietzsch, T., et al. 2012. Fiji: an open-source platform for biological-image analysis. Nat. Methods 9: 676–682. Nature Publishing Group.

Sexton, J.P., McIntyre, P.J., Angert, A.L. & Rice, K.J. 2009. Evolution and Ecology of Species Range Limits. Annu. Rev. Ecol. Evol. Syst. 40: 415–436. Annual Reviews.

Shu, Y., Du, Y. & Wang, J. 2011. Molecular characterization and expression patterns of Spodoptera litura heat shock protein 70/90, and their response to zinc stress. Comp. Biochem. Physiol. A. Mol. Integr. Physiol. 158: 102–10.

Sih, A. 2011. Effects of early stress on behavioral syndromes: An integrated adaptive perspective. Neurosci. Biobehav. Rev. 35: 1452–1465. Pergamon.

Slos, S. & Stoks, R. 2008. Predation risk induces stress proteins and reduces antioxidant defense. Funct. Ecol. 22: 637–642. Wiley/Blackwell (10.1111).

Sørensen, J.G., Kristensen, T.N. & Loeschcke, V. 2003. The evolutionary and ecological role of heat shock proteins. Ecol. Lett. 6: 1025–1037. Wiley/Blackwell (10.1111).

Sørensen, J.G. & Loeschcke, V. 2001. Larval crowding in Drosophila melanogaster induces Hsp70 expression, and leads to increased adult longevity and adult thermal stress resistance. J. Insect Physiol. 47: 1301–1307.

Stoks, R. 2001. Food stress and predator-induced stress shape developmental performance in a damselfly. Oecologia 127: 222–229. Springer Berlin Heidelberg.

Sunday, J.M., Bates, A.E. & Dulvy, N.K. 2012. Thermal tolerance and the global redistribution of animals. Nat. Clim. Chang. 2: 686–690. Nature Publishing Group.

Svensson, E.I., Abbott, J. & Hardling, R. 2005. Female polymorphism, frequency dependence, and rapid evolutionary dynamics in natural populations. Am. Nat. 165: 567–76. The University of Chicago Press.

Takahashi, Y., Kagawa, K., Svensson, E.I. & Kawata, M. 2014. Evolution of increased phenotypic diversity enhances population performance by reducing sexual harassment in damselflies. Nat. Commun. 5. Nature Research.

Van Gossum, H., Stoks, R. & De Bruyn, L. 2001. Frequency-dependent male mate harassment and intra-specific variation in its avoidance by females of the damselfly Ischnura elegans. Behav. Ecol. Sociobiol. 51: 69–75. Springer-Verlag.

Verberk, W.C.E.P. & Calosi, P. 2012. Oxygen limits heat tolerance and drives heat hardening in the aquatic nymphs of the gill breathing damselfly Calopteryx virgo (Linnaeus, 1758). J. Therm. Biol. 37: 224–229. Elsevier.

Wiens, J.J. 2011. The niche, biogeography and species interactions. Philos. Trans. R. Soc. Lond. B. Biol. Sci. 366: 2336–2350.

Wilson, R.S. & Franklin, C.E. 2002. Testing the beneficial acclimation hypothesis. Trends Ecol. Evol. 17: 66–70. Elsevier Current Trends.

Zhao, L. & Jones, W.A. 2012. Expression of heat shock proteins in insect stress responses. Invertebr. Surviv. J. 9: 93–101.

